# Discovery of unique loci that underlie nematode responses to benzimidazoles

**DOI:** 10.1101/116970

**Authors:** Mostafa Zamanian, Daniel E. Cook, Stefan Zdraljevic, Shannon C. Brady, Daehan Lee, Junho Lee, Erik C. Andersen

## Abstract

Parasitic nematodes impose a debilitating health and economic burden across much of the world. Nematode resistance to anthelmintic drugs threatens parasite control efforts in both human and veterinary medicine. Despite this threat, the genetic landscape of potential resistance mechanisms to these critical drugs remains largely unexplored. Here, we exploit natural variation in the model nematodes *Caenorhabditis elegans* and *Caenorhabditis briggsae* to discover quantitative trait loci (QTL) that control sensitivity to benzimidazoles widely used in human and animal medicine. High-throughput phenotyping of albendazole, fenbendazole, mebendazole, and thiabendazole responses in panels of recombinant lines led to the discovery of over 15 QTL in *C. elegans* and four QTL in *C. briggsae* associated with divergent responses to these anthelmintics. Many of these QTL are conserved across benzimidazole derivatives, but others show drug and dose specificity. We used near-isogenic lines to recapitulate and narrow the *C. elegans* albendazole QTL of largest effect and identified candidate variants correlated with the resistance phenotype. These QTL do not overlap with known benzimidazole resistance genes from parasitic nematodes and present specific new leads for the discovery of novel mechanisms of nematode benzimidazole resistance. Analyses of orthologous genes reveal significant conservation of candidate benzimidazole resistance genes in medically important parasitic nematodes. These data provide a basis for extending these approaches to other anthelmintic drug classes and a pathway towards validating new markers for anthelmintic resistance that can be deployed to improve parasite disease control.

**Author Summary:** The treatment of roundworm (nematode) infections in both humans and animals relies on a small number of anti-parasitic drugs. Resistance to these drugs has appeared in veterinary parasite populations and is a growing concern in human medicine. A better understanding of the genetic basis for parasite drug resistance can be used to help maintain the effectiveness of anti-parasitic drugs and to slow or to prevent the spread of drug resistance in parasite populations. This goal is hampered by the experimental intractability of nematode parasites. Here, we use non-parasitic model nematodes to systematically explore responses to the critical benzimidazole class of anti-parasitic compounds. Using a quantitative genetics approach, we discovered unique genomic intervals that control drug effects, and we identified differences in the genetic architectures of drug responses across compounds and doses. We were able to narrow a major-effect genomic region associated with albendazole resistance and to establish that candidate genes discovered in our genetic mappings are largely conserved in important human and animal parasites. This work provides new leads for understanding parasite drug resistance and contributes a powerful template that can be extended to other anti-parasitic drug classes.

## Introduction

Parasitic nematodes impose a debilitating health and economic burden across much of the developing world, conservatively resulting in the loss of 14 million disability-adjusted life years per annum^1^. This disease burden is partly curtailed by mass drug administration (MDA) programs that depend on the continued efficacy of a limited portfolio of anthelmintic drugs. Benzimidazoles are an indispensable component of this limited chemotherapeutic arsenal and three benzimidazole derivatives are considered ‘Essential Medicines’ by the World Health Organization. In veterinary medicine, aggressive benzimidazole chemotherapy has generated geographically widespread benzimidazole resistance^2^. The ongoing expansion of anthelmintic coverage in humans threatens a similar outcome^3’4^. Reduced cure rates suggestive of benzimidazole resistance have been reported for human soil-transmitted helminths, including the etiological agents of hookworm infection, trichuriasis, and ascariasis^5–9^. The sustainability of chemotherapy-based parasite control is jeopardized by our deficient knowledge of potential mechanisms of anthelmintic resistance and a paucity of molecular markers to detect and slow the spread of resistance alleles in parasite populations.

Traditional approaches to identify anthelmintic resistance markers rely on surveying candidate genes for polymorphisms^10^. Known mechanisms of nematode benzimidazole resistance have been limited to variants in the drug target beta-tubulin^10–13^. However, genetic differences in beta-tubulin genes do not explain all interspecific and intraspecific variation observed in benzimidazole efficacy^14^ or in responses to different benzimidazole derivative^15,16^. A more complete understanding of potential pathways to parasite benzimidazole resistance is necessary to help discover loci that are predictive of drug response. Although *C. elegans* has been historically indispensable to the discovery of mechanisms of action for benzimidazoles and other anthelmintics^17^, genetic variation within *C. elegans* has only recently been exploited to study phenotypic variation in anthelmintic responses^18^.

We used standing genetic variation and high-throughput quantitative phenotyping in two experimentally tractable *Caenorhabditis* species, *C. elegans* and *C. briggsae*, to identify genomic loci that control susceptibility to four clinically relevant benzimidazoles. Our findings reveal both conserved and drug-specific loci in each species that contribute to the effects of benzimidazoles on animal offspring production and growth rate. The genetic architectures of benzimidazole sensitivity and the specific genomic loci identified in this work provide new leads to identify genetic markers and molecular mechanisms that govern anthelmintic resistance in parasitic nematodes. We expect that translation of these leads will help to improve the detection and management of drug resistance in parasite populations.

## Methods

### High-throughput phenotyping assay

Strains were propagated for four generations to reduce transgenerational effects of starvation and bleach-synchronized before transfer to 96-well growth plates (~ 1 embryo/*μ*l in K medium). Hatched L1 larvae were fed HB101 bacterial lysate (5 mg/ml) and incubated for 48 hours at 20°C. L4 larvae were sorted into 96-well drug plates (three animals/well) using the COPAS BIOSORT large particle sorter (Union Biometrica). Drug plates contained anthelmintics dissolved in K medium at the desired final concentrations along with 1% DMSO, 10 mg/ml HB101 bacterial lysate, and 31.25 *μ*M kanamycin. These cultures were incubated for 96 hours at 20°C to allow development to the adult stage and the maturation of deposited embryos. Animals were fed a solution of 1 mg/ml bacterial lysate and 0.01 μM red fluorescent microspheres (Polysciences, cat. 19507-5) for five minutes prior to scoring. Animals were immobilized with 50 mM sodium azide, and the COPAS BIOSORT large particle sorter was used to measure a range of animal fitness traits including length, pharyngeal pumping (red fluorescence), and brood size^19,20^.

### Trait generation for dose responses and linkage mapping

Raw phenotype data collected from the COPAS BIOSORT large particle sorter were processed with the R package easysorter^21^. The function `read_data()` was used to distinguish animals from bubbles using a support vector machine (SVM). The functions `remove_contamination()` and `sumplate()` were used to mask contaminated wells and to calculate summary statistics across measured parameters. Parameters included time-of-flight (animal length), extinction (optical density), fluorescence (pharyngeal pumping), and total object count (brood size). Summary statistics included the mean and quantiles (10th, 25th, 50th, 75th, and 90th) for each of these parameters. Brood size was normalized to the sorted number of animals per well (n), while fluorescence was normalized to animal length (TOF). The `regress(assay=TRUE)` function was used to fit a linear model to account for differences in assays carried out on different days. Outliers were defined as observations that fall outside the IQR by at least twice the IQR and that do not group with at least 5% of the observations. Outliers were removed using the `bamf_prune()` function and `regress(assay=FALSE)` was used to fit a linear model (phenotype ~ control phenotype) to calculate drug effects with respect to control (DMSO solvent) conditions.

### Dose responses and selection of heritable doses

The dose-dependent phenotypic effects of benzimidazoles were assayed in technical quadruplicate across four genetically diverged *C. elegans* (N2, CB4856, DL238, and JU258) and *C. briggsae* (AF16, HK104, VT847, andED3035) strains. Phenotypes were measured using the high-throughput assay and trait generation pipeline described previously. Drug concentrations for linkage mapping experiments were selected based on broad-sense heritability calculations for traits of interest and with the goal of maximizing differences in sublethal drug effects between the parental strains used to generate recombinant lines (*C. elegans:* N2 and CB4856; *C. briggsae:* AF16 and HK104).

### Linkage mapping of fitness traits

Benzimidazole exposure phenotypes were measured for a population of 292 unique *C. elegans* recombinant inbred advanced intercross lines (RIAILs) resulting from an advanced intercross of N2 and CB4856, as well as 153 unique *C. briggsae* recombinant inbred lines (RILs) created using AF16 and HK104. These phenotypic data were collected and processed as described above. R/qtl^22^ was used to carry out marker regression on 1454 *C. elegans* markers and 1031 *C. briggsae* markers. QTL were detected by calculating logarithm of odds (LOD) scores for each marker and each trait as *−n(ln*(1 − *r*^2^)/2*l*n(10)), where *r* is the Pearson correlation coefficient between RIAIL genotypes at the marker and phenotype trait values^23^. Significance thresholds for QTL detection were calculated using 1000 permutations and a genome-wide error rate of 0.05. The marker with the maximal LOD score exceeding significance was retained as the peak QTL marker for each of three mapping iterations. QTL confidence intervals were defined by a 1.5 LOD drop from peak QTL markers. Broad-sense heritability was calculated using repeat measures of parental and recombinant strain phenotypes, as described previously^24^.

### Generation of near-isogenic lines

NILs were generated by backcrossing N2xCB4856 RIAILs to either parental strain for six generations, followed by six generations of selfing to homozygose the genome. Primers were optimized to genotype N2xCB4856 insertion-deletion variants immediately flanking introgression regions of interest. NIL reagents, primers, and PCR conditions are detailed in S1 Methods.

### Mutant strains

Existing *C. elegans* mutants *(alg-4(tm1184); alg-3(tm1155), ergo-1(tm1860), prg-1(n4357), ben-1(tm234))* were propagated alongside N2 and CB4856 to assay albendazole-response phenotypes. The *prg-1* mutant strain was backcrossed to N2 for 10 generations. Independent *prg-1* deletion strains were generated by CRISPR/Cas9-mediated gene editing using Cas9 ribonucleoprotein^25^. A co-CRISPR strategy targeting *dpy-10* was used to improve efficiency of screening for edits^26^. All Cas9 reagents were purchased through IDT (Skokie, IL). Alt-R tracrRNA (IDT, 1072532), *dpy-10* crRNA, and each *prg-1* crRNA (oECA2002 and oECA2003) were combined and incubated at 95°C for five minutes. Cas9 nuclease (IDT, 1074181) was added and the mix was incubated at room temperature for five minutes. Finally, the *dpy-10* repair template was added, and the final volume was brought to 5 *μ*L with nuclease-free water. The final concentrations are as follows: tracrRNA 13.6 *μ*M, *dpy-10* crRNA 4 *μ*M, each *prg-1* crRNA 9.6 *μ*M, Cas9 23.8 *μ*M, and *dpy-10* repair construct 1.34 *μ*M. The mix was centrifuged at maximum speed for five minutes, mouth pipetted into a pulled injection needle (World Precision Instruments, 1B100F-4), and injected into young adult N2 and CB4856 animals. Each injected animal was placed onto an individual 6 cm NGM plate approximately 18 hours post-injection. Roller F1 animals were placed onto individual 6 cm NGM plates when they reached the L4 stage, or when the Rol phenotype was apparent, and allowed to lay embryos. These F1 animals were then screened for large deletion events with PCR using primers oECA2004 and oECA2042. Non-Dpy, non-Rol progeny from edited F1 animals were propagated until homozygous and verified with Sanger sequencing. Primers and other reagents used to genotype backcross progeny and to generate CRISPR alleles are outlined in S1 Methods.

### Mutant and NIL strain phenotyping assays

NIL and mutant phenotyping assays were carried out with the high-throughput pipeline described above with at least two independent biological replicates carried out for each strain panel.

### Statistical analyses

Phenotype data are shown as Tukey box plots. Analyses were performed using R by one or two-tailed t-test (for two groups and specific hypotheses about direction of effect) or one-way ANOVA with Tukey’s multiple comparison test (for more than two groups). P-values less than 0.05 were considered significant. P-values for all statistical tests are provided in S9 Table.

### Gene interval annotation and parasite orthology analysis

*C. elegans* variants distinguishing N2 and CB4856^27^ were used to annotate QTL intervals with respect to existing gene annotations. *C. briggsae* variants distinguish AF16 and HK104 as well as their estimated functional consequences were produced from genomic sequence and annotation data using SnpEff^28^. For all *C. elegans* QTL identified in this study, parasite orthologs of genes with variants predicted to be of ’moderate’ or ’high’ impact were extracted with custom Python scripts. *Caenorhabditis* orthologs from the clade IV gastro-intestinal parasite *Stronygloides ratti* and the clade III human filarial parasite *Brugia malayi* were extracted from WormBase^29^.

### RNA-Seq pipeline

*C. elegans* strains N2 and CB4856 were bleach-synchronized and grown at 20°C for isolation of total RNA from young adult animals (60 hours post-embryo plating) using a liquid N2 freeze-cracking protocol with TRIzol (Life Technologies). RNA was collected from four independent biological replicates per strain. Sample RNA concentration and quality were assessed via Agilent Bioanalyzer. mRNA libraries were prepared from RNA samples using the TruSeq Stranded mRNA Library Prep Kit with oligo-dT selection (Illumina). All samples were sequenced using the Illumina HiSeq 2500 platform with a single-end 50 bp read setting (University of Chicago Genomics Facility) and demultiplexed for downstream analyses. Mean per-sample yield was 10.0 Gb for mRNA-seq samples. Reads were adapter and quality trimmed using Trimmomatic^30^. HiSAT2 and StringTie^31^ were used to produce raw and TPM (transcripts per million) read counts for annotated genes. DESeq2^32^ was used to identify differentially expressed genes. The complete RNA-seq pipeline was implemented with Nextflow^33^ and is publicly available through GitHub (github.com/AndersenLab/BZRNA-seq-nf).

### Data Availability

Linkage mapping and phenotype data and all scripts used in the analyses of these and any other data are available through GitHub (github.com/AndersenLab/C.-elegans-Benzimidazole-Resistance-Manuscript/). RNA sequencing data are available in the NCBI SRA under accession number [NUMBER].

## Results and Discussion

### Natural variation in *C. elegans* responses to benzimidazole anthelmintics

We examined natural variation in *C. elegans* responses to four benzimidazoles (albendazole, fenbendazole, mebendazole, and thiabendazole) that are widely used in human and veterinary medicine. Dose responses were performed on a set of four genetically diverged strains using a flow-based, large-particle analysis device (COPAS Biosort, Union Biometrica) for high-throughput quantification of anthelmintic effects on animal fitness traits, including offspring production and growth rate. This platform enables trait measurements at unprecedented scales in a metazoan system^19^ (S1 Fig.). In brief, strains are synchronized and dispensed into 96-well microtiter plates containing anthelmintic drugs or no drug (DMSO). Conditions are strictly controlled for temperature, humidity, food source, and culture mixing. The COPAS Biosort measures the length, optical density, and fluorescence (green, yellow, and red) of every nematode from populations grown in 96-well microtiter plates. This high-throughput platform enables the measurement of animal size, fecundity, and feeding behavior. As *C. elegans* grows, animals get longer^34^, so length measurements are a proxy for developmental stage and growth rate. Fecundity is assessed by counting the number of offspring produced by a defined number of parent animals in each well. Feeding rate is quantified by exposing animals to fluorescent microspheres (Fluoresbrite Fluorescent Microspheres, Polysciences Inc.), which are the same size as bacterial food, and then measuring fluorescence after a defined period of time. Anthelmintic drugs reduce growth rate, offspring production, and muscle activity, so these traits are directly relevant to anthelmintic mechanisms of action in parasitic nematodes^35, 36^. Assay measurements with longer animals, more offspring, and/or more fluorescence indicate that that strain is more resistant to an anthelmintic than strains with shorter animals, fewer offspring, and/or less fluorescence. All drugs showed heritable dose-dependent effects for at least one of these major phenotypic categories (S2 Fig.). Drug concentrations that exhibited high broad-sense heritability and that maximized strain-specific differences were chosen for quantitative genetic mappings. To assess the potential effects of dose on the genetic architecture of drug sensitivity, two concentrations of fenbendazole and thiabendazole were selected. These sublethal concentrations fall within the range of previous studies in C. elegans^11,37^ and likely correspond to pharmacologically relevant drug accumulation levels in parasitic nematodes^38,39^.

### Discovery of quantitative trait loci (QTL) associated with benzimidazole responses in *C. elegans*

From our dose response assays of four genetically diverse *C. elegans* strains, we found that the laboratory strain from Bristol, England (strain N2) and a wild strain from Hawaii, U.S.A. (strain CB4856) differed in responses to benzimidazoles. We can use the natural differences between these two strains to identify the variants that contribute to divergent anthelmintic responses. Previously, these two strains were crossed to create a collection of recombinant inbred advanced intercross lines (RIAILs) to facilitate linkage mapping approaches^19,40^. Using 292 RIAILs, we identified 15 quantitative trait loci (QTL) that that each explain greater than 5% of trait variation in benzimidazole susceptibility (Figure 1A, S1 Table). Many of these QTL span multiple animal fitness traits (S3 Fig.). The trait groupings can be observed by the correlation structure of measured parameters and robust across summary statistics (mean, median, 75th quantile, and 90th quantile) for these parameters (S4 Fig.). Benzimidazole sensitivity involved the contributions of multiple loci for many drug-trait combinations. Within this complex trait landscape, we identified QTL that are common but also some that are unique across drugs and doses. Drug-specific QTL provide new leads to explain observed differences in clinical efficacy among benzimidazole compounds. By contrast, dose-specific QTL reveal the engagement of different genetic determinants of benzimidazole response as a function of drug exposure.

**Figure 1.**
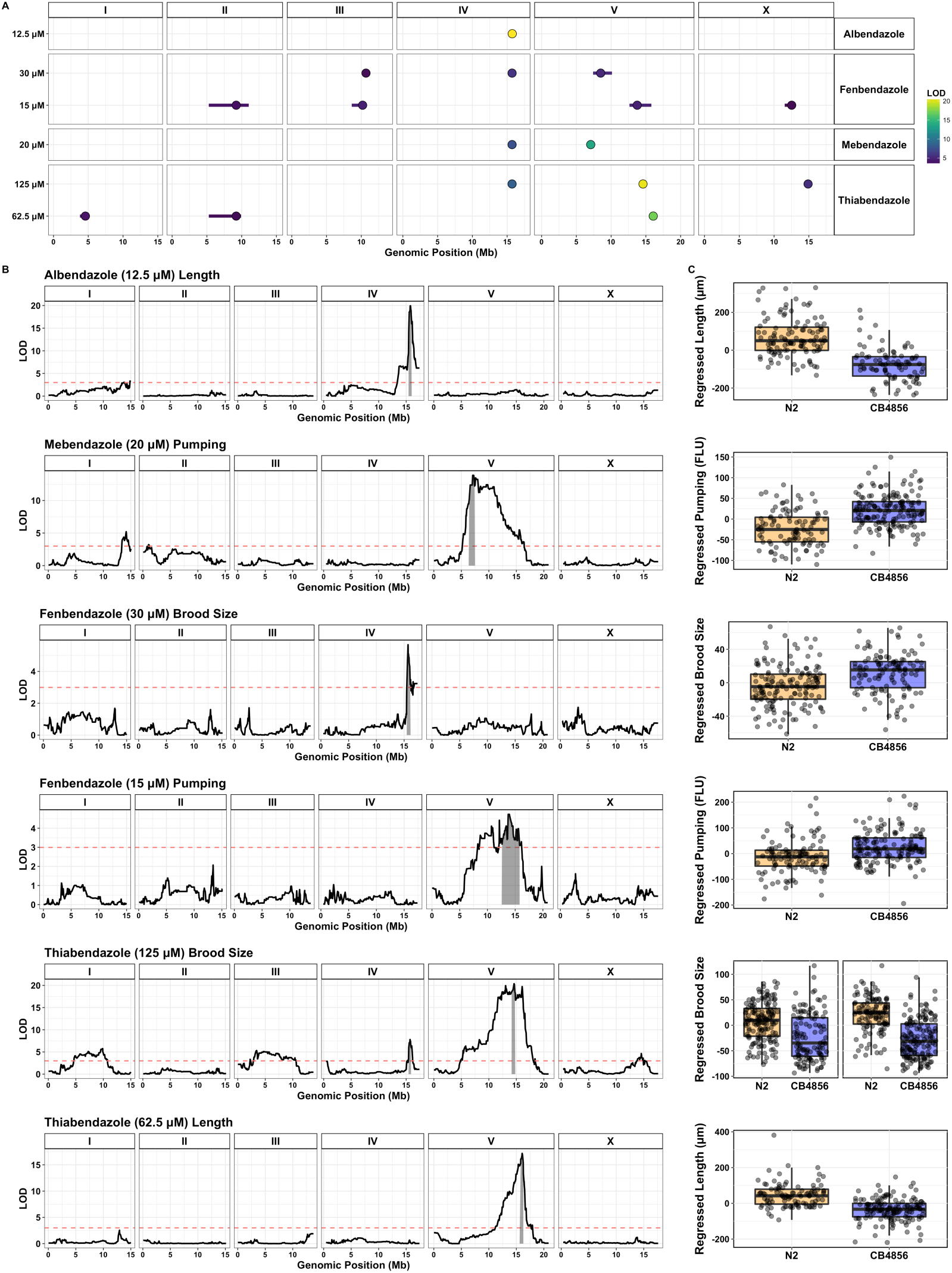
Discovery of benzimidazole response QTL in *C. elegans*. (A) Results of *C. elegans* linkage mapping experiments are shown for the five drug-dose conditions tested. QTL peak markers (circles) and confidence intervals (lines) are depicted. Fill color corresponds to the QTL significance (LOD) score. Overlapping QTL for a given condition are represented by the trait with the highest significance score. In total, 15 non-overlapping QTL were identified across these conditions. (B) Linkage mapping plots for the QTL of highest significance for each drug condition. The plots show genomic position on the x-axis and significance (maximum LOD score across mapping iterations) on the y-axis. QTL confidence intervals (e.g., chromosome IV: 15.47 - 15.91 Mb for 12.5 *μ*M albendazole) are shaded in gray and the red dashed line represents the genome-wide significance threshold for the first mapping iteration (LOD score for 12.5 *μ*M albendazole QTL peak marker = 19.93; LOD threshold = 2.99). (C) Tukey box plots showing the phenotypic split of RIAILs that have either the N2 or CB4856 genotype at the peak QTL marker position(s).

We discovered a major QTL associated with the effects of albendazole on animal size traits. This QTL is localized to a 441 kb interval on chromosome IV (15.47 - 15.91 Mb) and explains 36% of albendazole-induced variation in animal length among the strains (Figure 1B). This major-effect albendazole QTL extends to pharyngeal pumping and is also associated with small differences in brood size (S3 Fig.). An overlapping QTL associated with animal size and pumping behavior was identified for 30 *μ*M fenbendazole and 20 *μ*M mebendazole. A distinct size and pumping-associated QTL was mapped for 20 *μ*M mebendazole on the left arm of chromosome V that does not overlap with loci identified for other benzimidazole derivatives. Another highly significant QTL on chromosome V (14.15 - 14.72 Mb) explains 29% of thiabendazole (125 *μ*M)-induced variation in brood size between the strains. This QTL extends to length and pharyngeal pumping traits for both 62.5 *μ*M and 125 *μ*M thiabendazole, and partly overlaps with a QTL associated with differences in pumping behavior in 15 *μ*M fenbendazole. A number of unique loci with smaller effect sizes were mapped for fenbendazole sensitivity at 15 and 30 *μ*M, with only one overlapping QTL apparent across the two drug concentrations. Plots of phenotype as a function of peak QTL marker genotype (Figure 1C) show that the direction of effect varies across drugs and traits, indicating that both N2 and CB4856 strains possess alleles contributing to both benzimidazole sensitivity and resistance.

### Near-isogenic lines recapitulate and narrow major-effect albendazole QTL

The albendazole QTL confidence interval contains 17 protein-coding genes with coding variants and falls within the large Piwi-interacting RNA (piRNA) cluster on chromosome IV (13.5-17.2 Mbs)^41^. In the linkage mapping experiment, recombinant strains with the N2 genotype at this chromosome IV QTL locus (15.47 - 15.91 Mb) were more resistant to incubation in 12.5 *μ*M albendazole than those strains with the CB4856 genotype, with CB4856 animals exhibiting a significantly greater decrease in length (Figure 1C, top). To narrow this interval to a smaller region, we measured the albendazole response phenotypes of near-isogenic lines (NILs) with QTL genomic regions derived from either the N2 or CB4856 strains introgressed into the opposite genetic background (Figure 2A). NILs with the N2 genotype spanning the 15.57 - 15.65 Mb interval of chromosome IV exhibited greater resistance to albendazole compared to the parental CB4856 strain and, conversely, a NIL with the CB4856 genotype in this region was more sensitive to albendazole when compared to the parental N2 strain (Figure 2A, bottom). These results are consistent with the QTL direction of effect, and we expect that the variant(s) of largest effect fall within this narrowed interval. Significant differences among NILs within the resistance and sensitive groupings suggest that additional variants and epistatic interactions within the complete QTL interval influence albendazole susceptibility.

**Figure 2.**
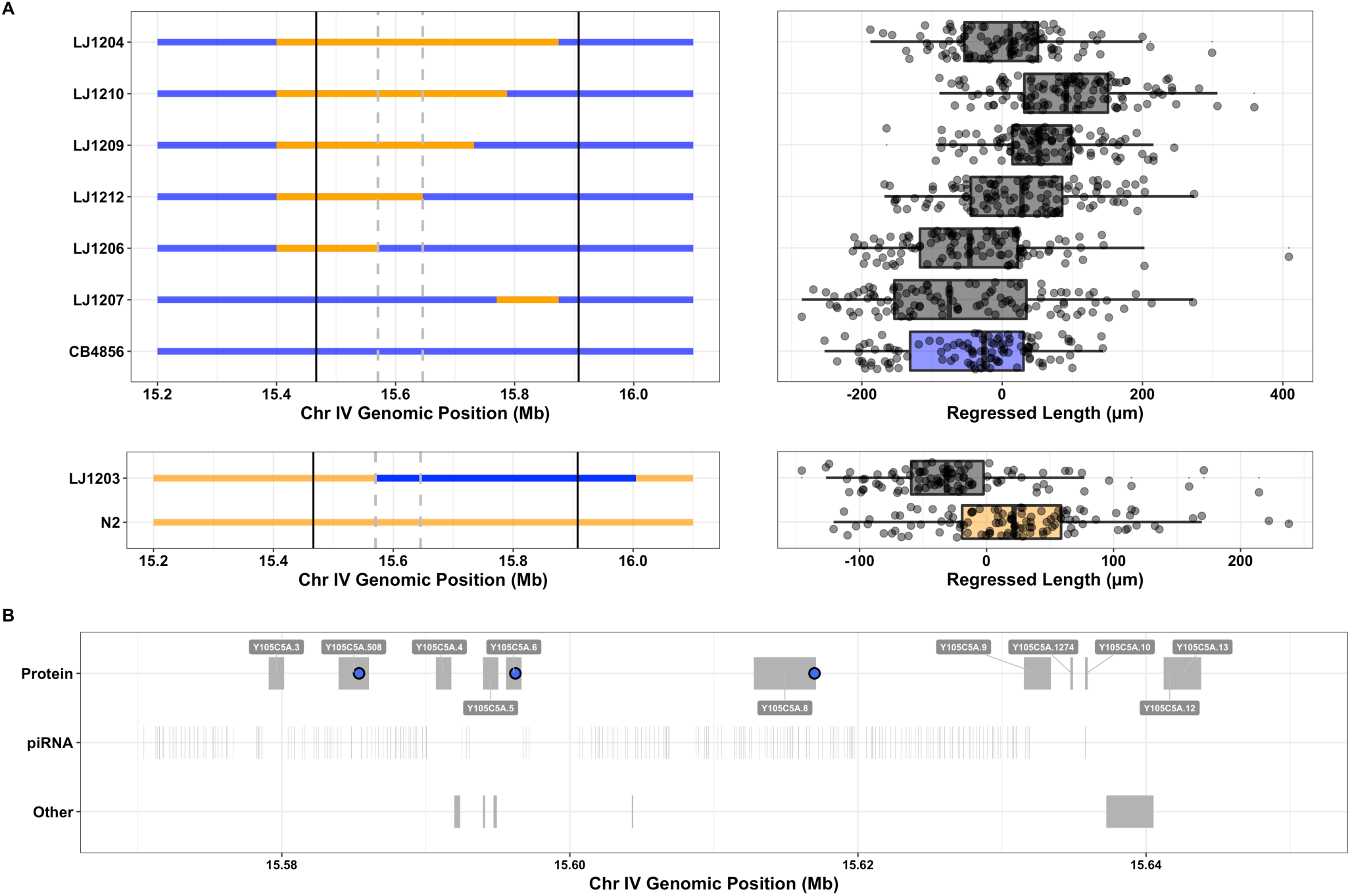
Narrowing of the albendazole QTL interval using near-isogenic lines. NIL genotypes (A) and corresponding Tukey box plots of NIL phenotypes (B) are shown with respect to their parental strain. NILs constructed via introgression of N2 into the CB4586 background can be classified into albendazole-sensitive (LJ1206 and LJ1207) and resistant groups (LJ1204, LJ1209, LJ1210, and LJ1212). This phenotypic separation allows for the tentative narrowing of the QTL interval (chromosome IV: 15.47 - 15.91 Mb) to a much smaller region (chromosome IV: 15.57 - 15.65 Mb) that contains one or more causative variants. Solid and dashed-vertical lines are used to mark the original QTL interval and the NIL-narrowed interval, respectively. Statistically significant differences among NILs in the resistant group suggest that additional variants outside of this narrowed interval contribute to phenotypic variance through epistatic interactions. Comparison of the N2 parental strain and LJ1203 are consistent with at least one causal variant contained within the narrowed interval. ANOVA with post-hoc Tukey HSD test was used to compare NILs used in these assays (p-values reported in S9 Table). (C) Variants distinguishing N2 and CB4856 within the NIL-narrowed interval are highlighted with respect to gene annotations. Candidate albendazole resistance variants include nonsynonymous and splice-donor variants in protein-coding genes (blue circles). No variants occur within genes encoding many non-coding RNA biotypes (tRNAs, miRNAs, and snoRNAs), though at least 276 variants occur within annotated piRNAs within the narrowed interval (S3 Table, not plotted).

The NIL interval was annotated with known variants distinguishing N2 and CB4856 and their estimated functional consequences (Figure 2B, S2 Table, and S3 Table). The presence of large numbers of annotated piRNAs added to the complexity of analyzing genome variation in both the complete QTL interval and the NIL-narrowed interval. We considered protein-coding and non-coding RNA (ncRNA) variation as potentially causal to the albendazole-resistance phenotype. Only three proteins in the narrowed interval are predicted to have altered functions as a result of single nucleotide variants (SNVs). However, the plausibility of these specific candidate genes was dampened by a number of factors. *Y105C5A.508* is curated as a short and likely pseudogenic transcript, which shares no homology with proteins in species with available sequence data. Additionally, we did not detect consistent expression of this gene using RNA-seq in either the N2 or CB4856 strains. The gene *pqn-79* has a predicted coding variant (Thr194Ala) but belongs to a highly redundant protein family that has > 99.5% sequence identity with at least three other homologs. None of these homologs exhibits coding variation, but the possibility of a dominant-negative or gene dosage effect cannot be excluded. The gene *Y105C5A.8* codes for a protein of unknown function that is predicted to contain a splice-donor variant, but we found no differences in overall expression (S4 Table) or splice form abundance (S5 Fig) between N2 and CB4856 for this gene. We therefore originally hypothesized that albendazole resistance is more likely a function of variation in the non-coding RNA (ncRNA) complement, specifically the piRNA-encoding genes. 1,684 SNVs occur in known piRNAs in the complete QTL interval, and 276 of these fall within the NIL-narrowed interval (S3 Table).

To test this hypothesis, we generated NILs encompassing the broader piRNA-enriched region on chromosome IV (13.5 - 17.2 Mbs). These NIL strains robustly recapitulated the QTL direction of effect (S6A Fig.). Next, we analyzed the responses of small RNA pathway mutants to albendazole in an effort to perturb the large number of diverse piRNAs. 21U-RNAs/piRNAs are regulated by the Piwi Argonaute PRG-1^42^. We hypothesized that albendazole sensitivity should be dependent on *prg-1* function. We tested mutants in all three Argonaute genes that encode proteins that interact with primary small RNAs immediately upstream of WAGO-associated 22G RNA generation *(ergo-1, alg-4; alg-3,* and *prg-1)^43^.* ERGO-1 and ALG-3/4 engage 26G RNAs, while PRG-1 coordinates the processing of piRNAs. Of the mutations in these genes, a back-crossed *prg-1* N2 strain conferred greater albendazole sensitivity compared to N2 (S6B Fig.). Concerned with the relative sickness of this mutant strain in no drug (DMSO) control, we generated and tested two independent *prg-1* loss-of-function alleles in both the N2 and CB4865 backgrounds using CRISPR/Cas9 genome editing. These new *prg-1* knock-out strains were not significantly different from wild-type parents in the expected direction of effect across three replicate assays (S7 Fig.). Although we have not identified the causal genetic locus, it is possible that the factors underlying the albendazole resistance phenotype are a mixture of coding or regulatory variants that interact and affect gene function. Numerous expression QTL (eQTL) have also previously been mapped to the major-effect albendazole QTL region, potentially connecting candidate variants to *cis* or *trans* regulatory targets that affect drug sensitivity^44^ (S5 Table).

### Discovery of quantitative trait loci (QTL) associated with benzimidazole responses in *C. briggsae*

To compare benzimidazole-resistance loci across nematode species, we examined variation in the responses of *Caenorhabditis briggsae* strains to the same set of benzimidazole compounds that we studied in *C. elegans.* Dose responses and heritability calculations (S8 Fig. and S9 Fig.) were used to select concentrations for linkage mapping experiments. Linkage mapping was carried out with a collection of 153 recombinant inbred lines (RILs) created using the parental strains AF16 and HK104^45^, which led to the discovery of four QTL for the tested drugs (Figure 3A, S10 Fig., and S6 Table). The *C. briggsae* QTL of largest effect is found on the left arm of chromosome IV (2.56 - 3.45 Mb) and explains approximately 18% of fenbendazole-induced variation in fecundity between the parental strains (Figure 3B). QTL of smaller effect were identified on the center of chromosomes V for thiabendazole and on the left arms of chromosomes V and X for albendazole. No QTL were identified for mebendazole across the examined set of fitness traits.

**Figure 3.**
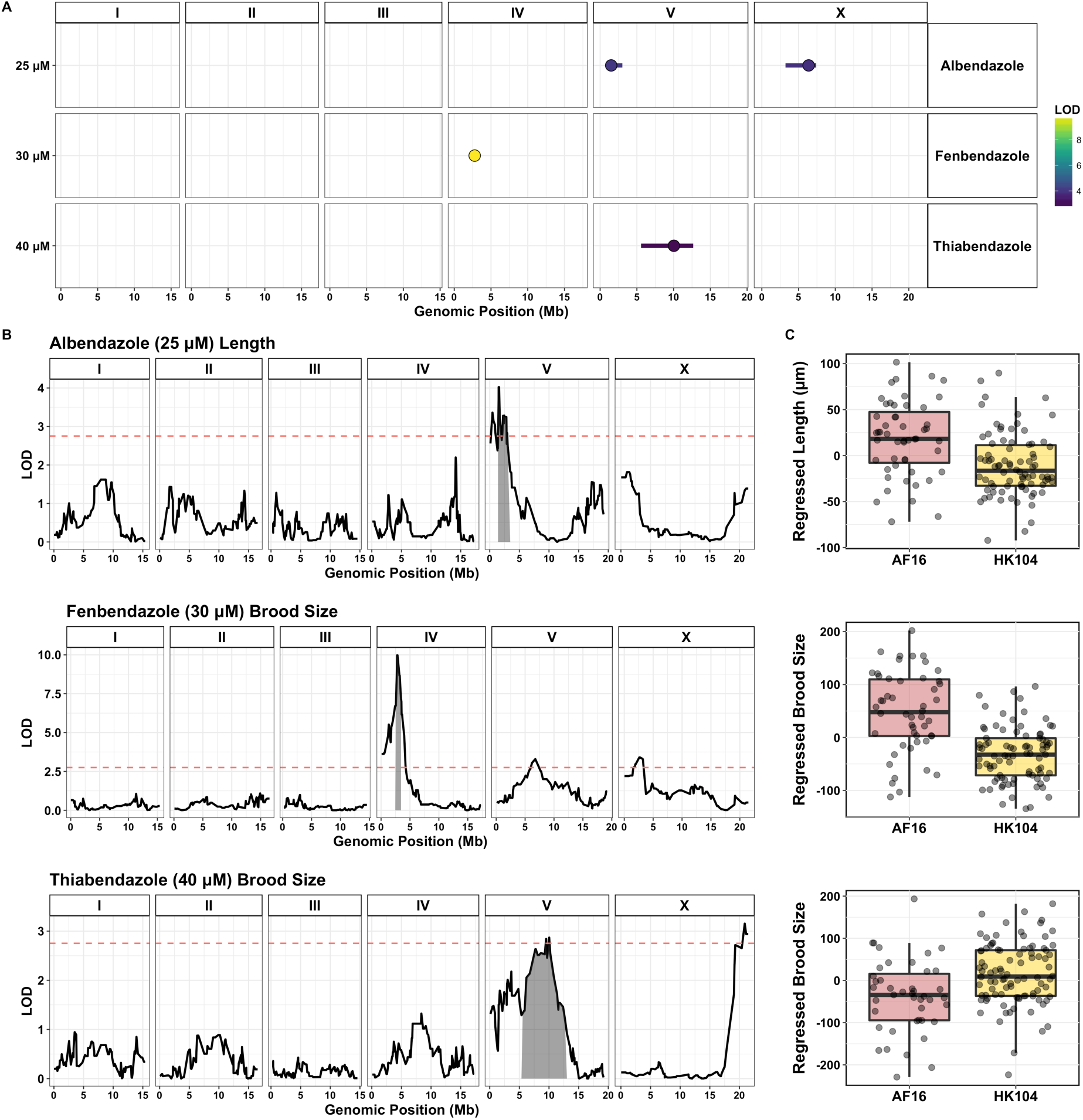
Discovery of benzimidazole response QTL in *C. briggsae*. (A) Results of *C. briggsae* linkage mapping experiments are shown for four tested drugs as grouped by trait category. QTL peak markers (circles) and confidence intervals (lines) are depicted. Fill color corresponds to the QTL LOD score. In total, four non-overlapping QTL were identified across conditions. (B) Linkage mapping plots are shown for the QTL of highest significance for each drug, including a major QTL associated with the effect of fenbendazole on brood size, and to a lesser extent, length and pumping traits. Plots show genomic position along the x-axis and significance (maximum LOD score across mapping iterations) along the y-axis. The QTL confidence intervals (chromosome IV: 2.56 - 3.45 Mb for 30 *μ*M fenbendazole) are shaded in gray and the red dashed line represents the genome-wide correction significance threshold for the first mapping iteration (LOD score for fenbendazole QTL peak marker = 9.97; LOD threshold = 2.75). The fenbendazole QTL explains 18% of trait variation (effect size = 0.70). (C) Tukey box plots showing the phenotypic split of RIAILs that have either the AF16 or HK104 genotype at the peak QTL marker positions. AF16 animals are significant more resistant to fenbendazole exposure than HK104 animals, exhibiting much larger brood sizes in the presence of drug.

Strikingly, the major fenbendazole QTL falls within the primary *C. briggsae* piRNA cluster (Chr IV: 0 - 6.9 Mb). Despite tens of millions of years of evolutionary distance^46^, the most significant *C. elegans* and *C. briggsae* benzimdazole QTL were found to occur in genomic regions that are syntenic between species^47^ (Figure 4A). The *C. briggsae* fenbendazole QTL region contains 84 protein-coding genes with variants (S7 Table), however, none of these genes are orthologous to variant-containing genes within the *C. elegans* albendazole and fenbedazole QTL intervals. This result suggests that different gene(s) and mechanisms account for the drug effects mapped to these syntenic loci across species. piRNA-encoding genes that densely cover this locus could potentially underlie the fenbendazole-resistance phenotype in *C. briggsae,* but no piRNAs are shared between *C. elegans* and *C. briggsae.*

**Figure 4.**
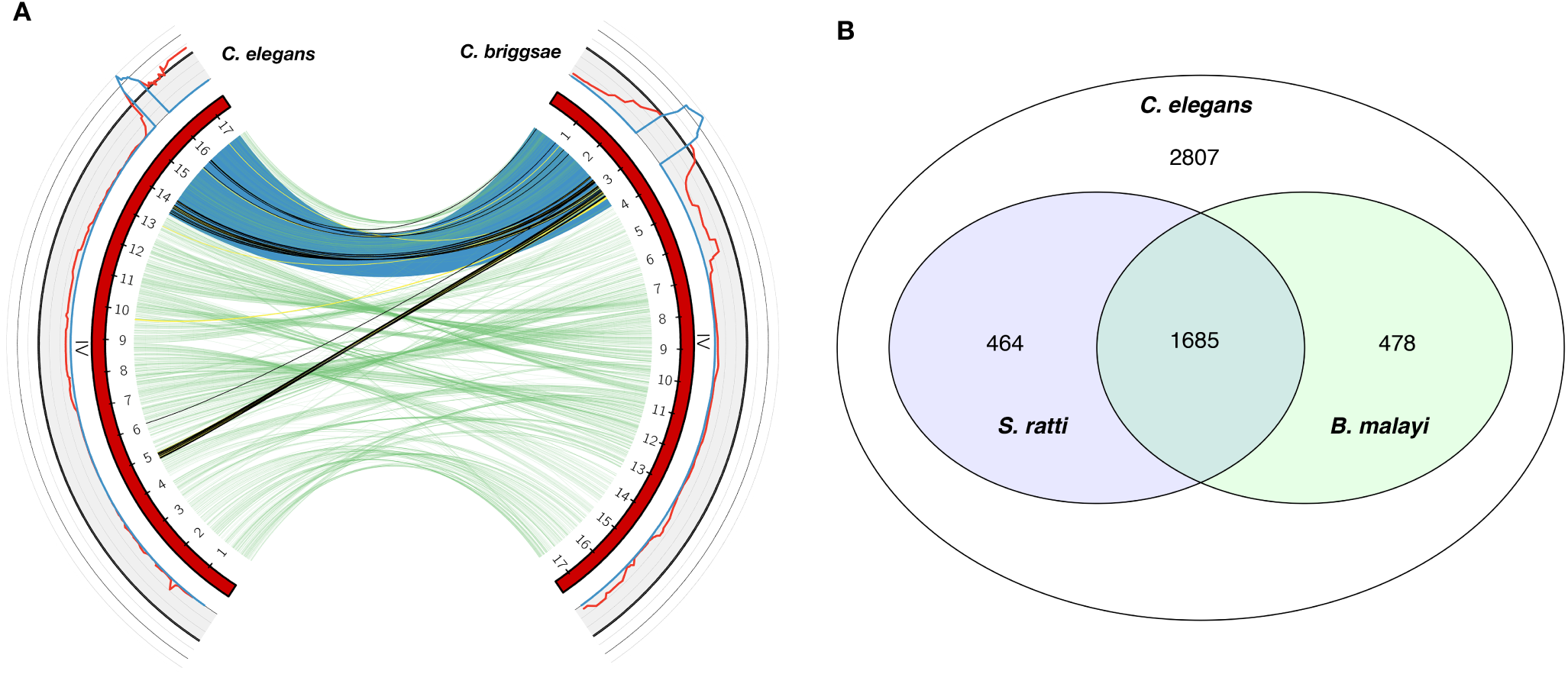
(A) Synteny between major-effect *C. elegans* albendazole QTL and *C. briggsae* fenbendazole QTL on chromosome IV. Circos^53^ plot showing synteny of orthologous gene pairs on chromosome IV. The outer track shows linkage plots for the *C. elegans* albendazole and *C. briggsae* fenbendazole mappings (red line: LOD scores; blue line: QTL confidence intervals; black line: significance thresholds). Both QTL intervals fall within major piRNA clusters in both species (piRNA cluster beginning and end sites shaded in blue). Inner green links represent all orthologs that occur outside of the QTL intervals, yellow links represent protein-coding orthologs with at least one member inside either QTL interval but with no detected variants (58 pairs), and black links represent orthologs with at least one member inside either QTL interval and with a detected variant (38 pairs). No orthologs were identified across species that both contained variants and fell within their respective QTL confidence intervals. This result suggests that different genes and mechanisms account for the drug effects mapped to these loci. (A) Conservation of candidate *C. elegans* benzimidazole resistance genes in the parasitic nematodes *S. ratti* and *B. malayi.* Venn diagram showing that a substantial proportion of the QTL-contained protein-coding genes with predicted functional variants (5,434 total) are conserved in representative clade IV and clade III parasites. Approximately 40% of the candidate resistance genes have identifiable orthologs in the clade III human filarial parasite *Brugia malayi* (2,163 of 5,434) and the clade IV model gastro-intestinal parasite *Strongyloides ratti* (2,149 of 5,434), and one-third are conserved across all three species (1,685 of 5,434).

### Candidate benzimidazole resistant genes are generally conserved in parasitic nematode species

To explore the potential translation of these loci to parasites, we looked at conservation of candidate *C. elegans* resistance genes(s) in parasitic nematodes. Specifically, we examined conservation of protein-coding genes that fall within benzimidazole QTL confidence intervals and contain predicted functional variants. Substantial fractions of these candidate genes have identifiable orthologs in representative parasites from other clades. Approximately 40% of the 5,434 candidate resistance genes across all identified QTL have orthologs in the clade III human filarial parasite *Brugia malayi* and the clade IV model gastro-intestinal parasite *Strongyloides ratti,* and 31% are conserved across all three species (Figure 4B and S8 Table). Despite a conservative threshold for assignment of orthology pairs, we found a high likelihood that gene(s) validated in this system as modulators of benzimidazole response have counterparts in parasite genomes. The narrowing of these QTL in *Caenorhabditis* species and the validation of the independent effects of genes on phenotypes has the potential to discover novel benzimidazole modes of action and resistance. Anthelmintic resistance gene variants validated in the genetically tractable *Caenorhabditis* species can be evaluated with respect to effect size and mutation type *(e.g.,* gain or loss-of-function). For genes with parasite orthologs, the effects of gene loss-of-function on parasite drug response can be assayed using targeted genetic perturbation techniques (*e.g.*, RNA interference) established in various helminth species. The expansion of the helminth genetic toolkit to CRISPR/Cas9 genome editing^48,49^ will pave the way for more precise mappings of mutations and effects. These *Caenorhabditis* data complement efforts to improve the resolution of parasite population genomics data^50^, as well as efforts to carry out genetics studies in helminth species where feasible^51,52^. More broadly, we expect that these approaches and data can hasten better models and markers for the development of anthelmintic resistance in human and animal parasite control.

## Funding

E.C.A. is a Pew Scholar in the Biomedical Sciences, supported by the Pew Charitable Trusts. Funding for E.C.A. is also provided by a Basil O’Connor Starter Research Award from the March of Dimes Foundation and an NIH NIAID grant (R21AI121836). Funding for M.Z. is provided by an NIH NIAID grant (K22AI125473). S.C.B. and S.Z. were supported by the Biotechnology Training Grant (T32GM008449) and the Cellular and Molecular Basis of Disease Training Grant (T32GM008061), respectively.

## Acknowledgements

The authors thank Sam Rosenberg and Robyn Tanny for help with the high-throughput phenotyping assay. The authors additionally thank Matteo Di Bernardo for help in NIL construction, as well as members of the Andersen laboratory for critical comments on the manuscript.

## Author contributions

M.Z. and E.C.A. designed the experiments, analyzed and interpreted the data, and wrote the manuscript. D.L. and J.L. provided reagents. M.Z. and D.E.C. analyzed genomic data. S.C.B. and S.Z. created the *prg-1* deletion alleles and scored BZ responses across NILs and mutants.

## Supporting Information: Figures

**S1 Fig.**
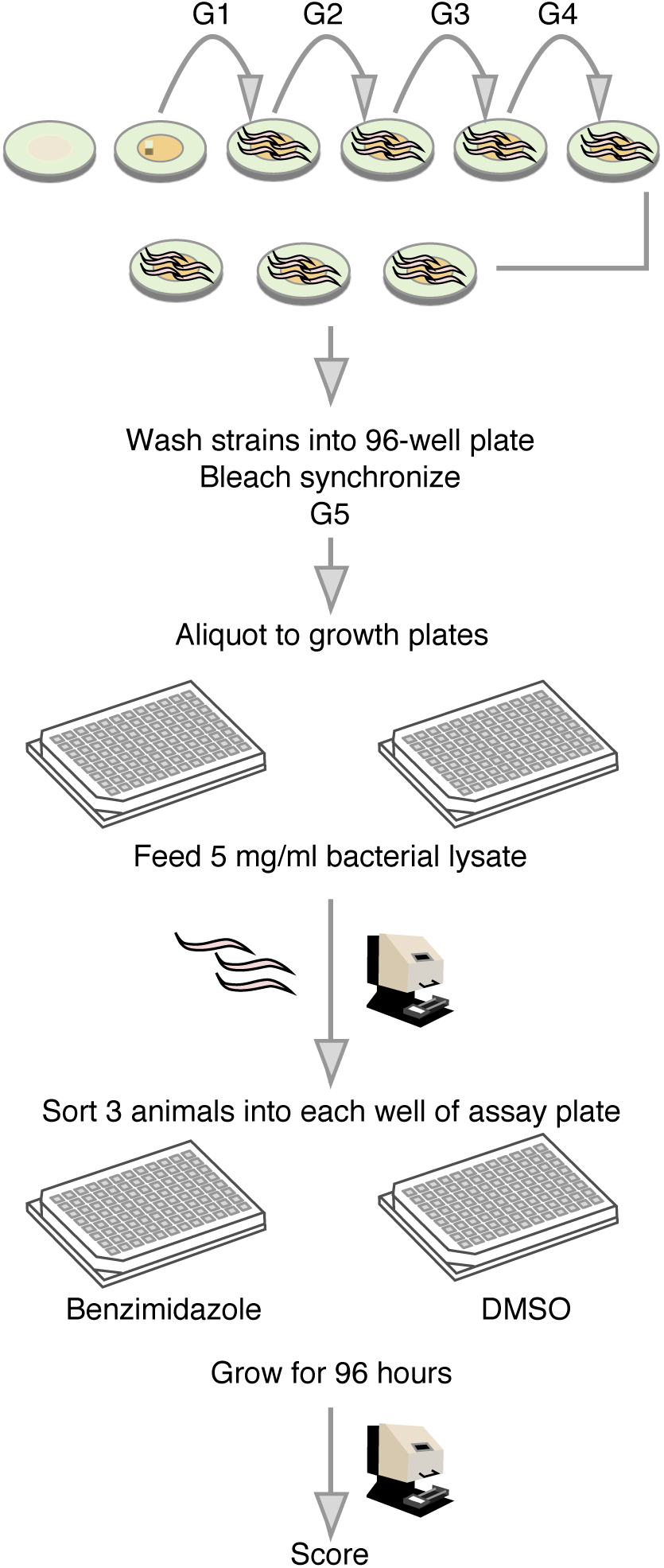
High-throughput sorter assay. Assay detailed in the Methods section.

**S2 Fig.**
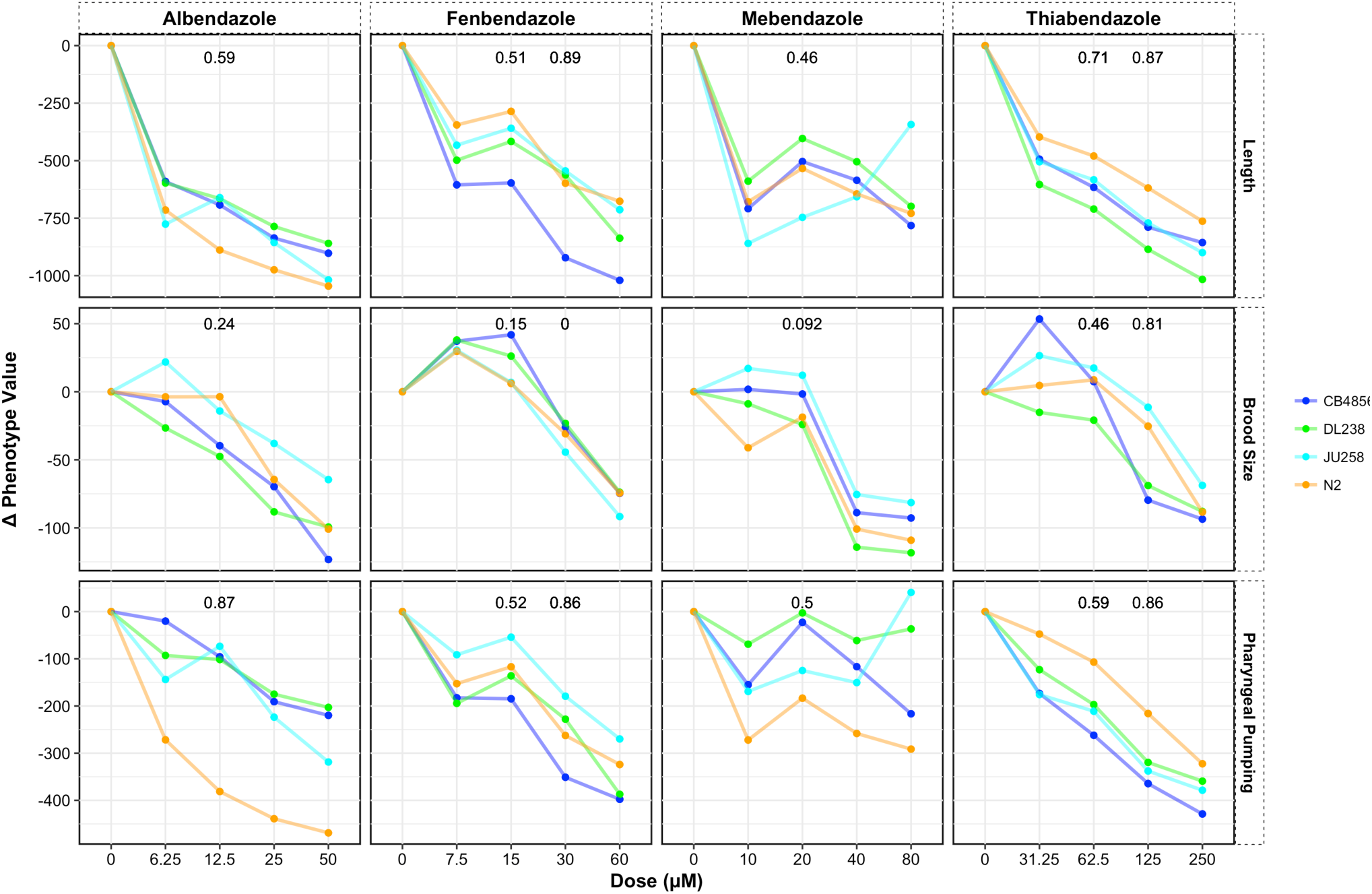
*C. elegans* benzimidazole dose responses and heritability calculations. Dose responses were carried out with four genetically diverged strains of *C. elegans.* Phenotypic responses to four drugs are shown with a representative trait for each primary trait group (length, brood size, and pharyngeal pumping). Heritability values are shown for doses used in subsequent linkage mapping experiments.

**S3 Fig.**
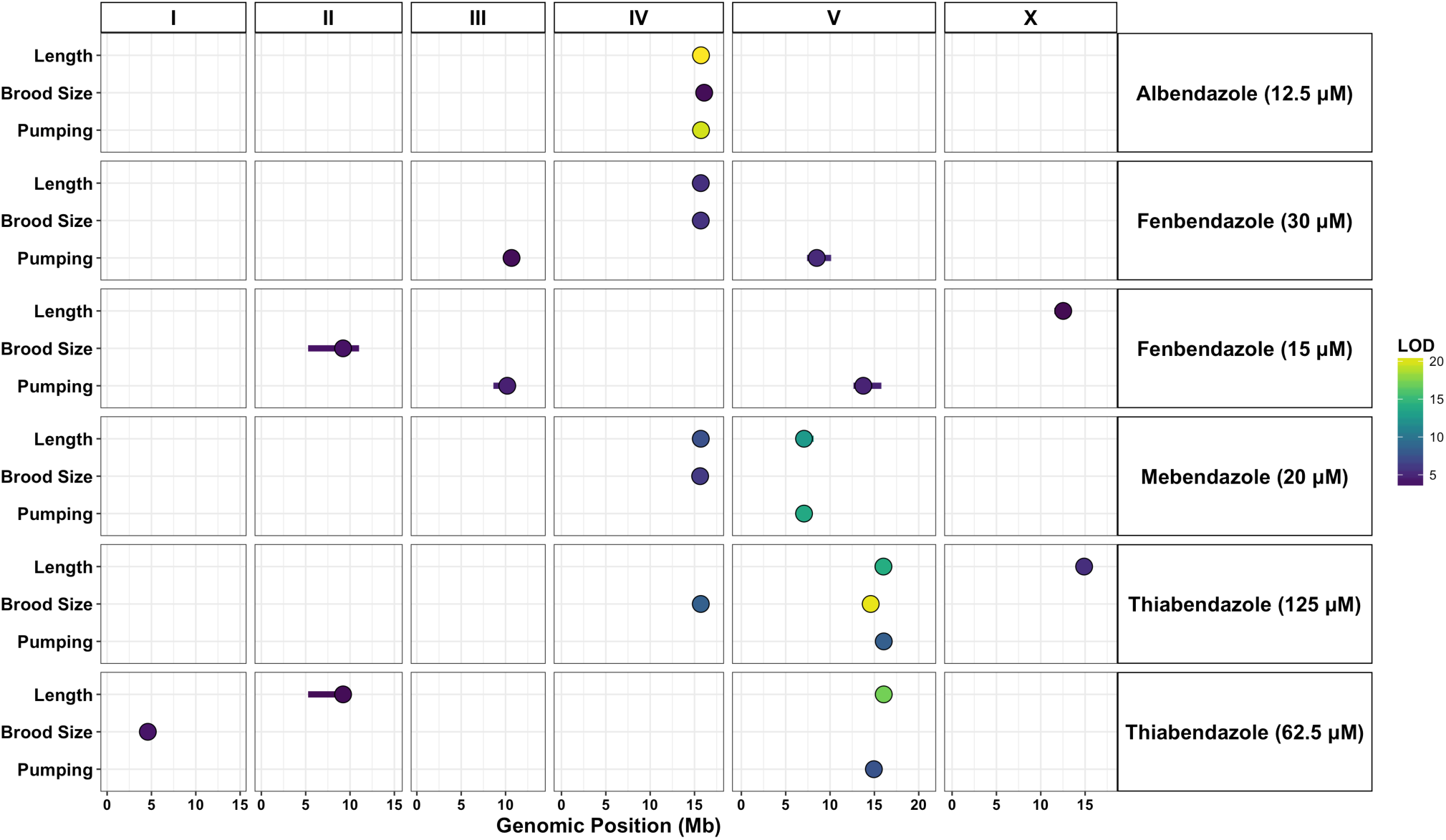
*C. elegans* benzimidazole QTL grouped by drug and trait. Results of *C. elegans* linkage mapping experiments are shown for the six drug-dose conditions tested and separated by correlated trait group. QTL peak markers (circles) and confidence intervals (lines) are depicted. Fill color corresponds to the QTL LOD score. Overlapping QTL for a given condition-trait group pair are represented by the trait with the highest significance score.

**S4 Fig.**
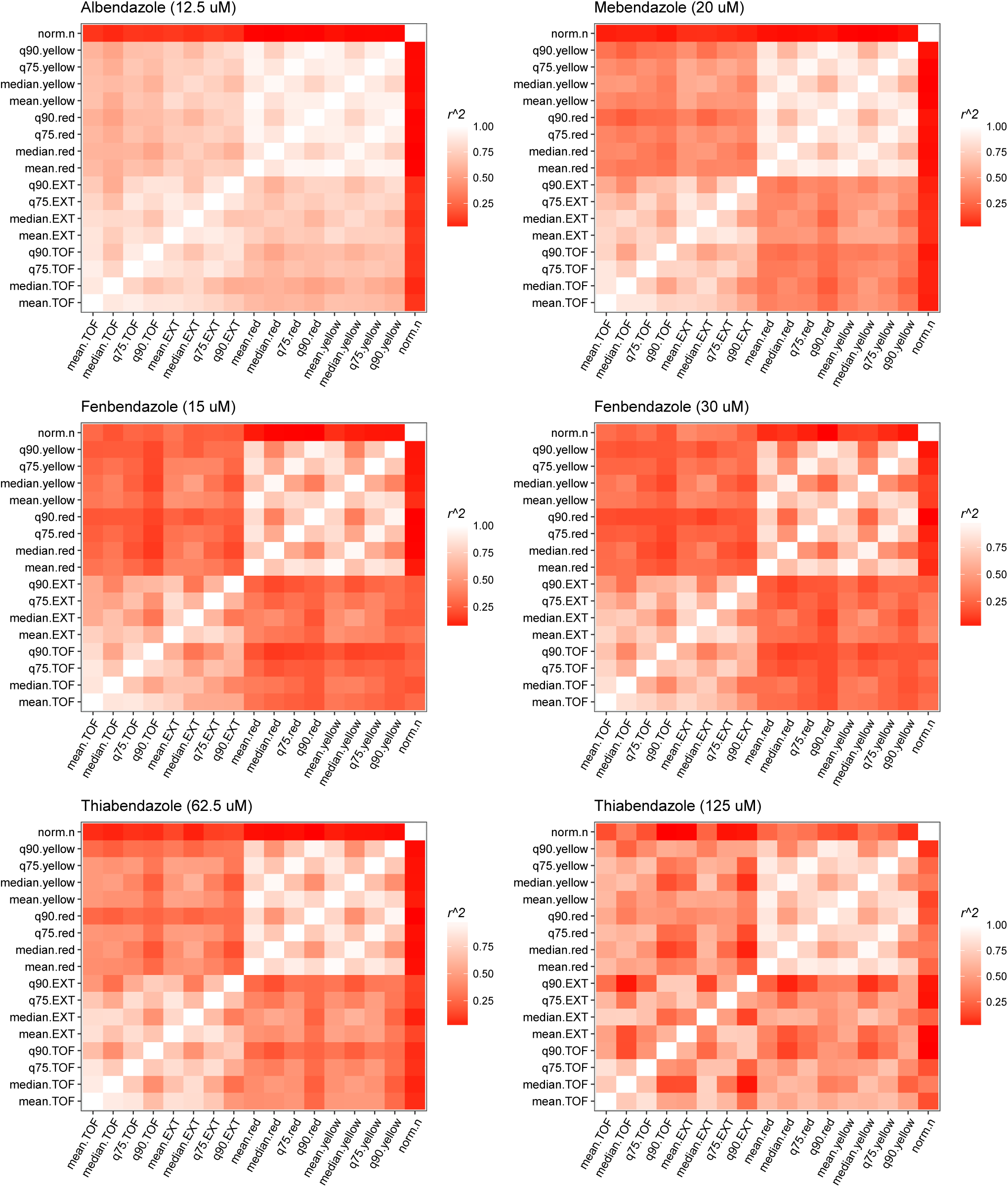
*C. elegans* summary statistic correlations for trait groupings. The correlation structure (Pearson’s correlation coefficient) of summary statistics for measured parameters of animal size (time-of-flight (TOF) and optical density (EXT)), pharyngeal pumping (red and yellow fluorescence), and brood size (norm.n) are shown for each drug and dose combination used in linkage mapping.

**S5 Fig.**
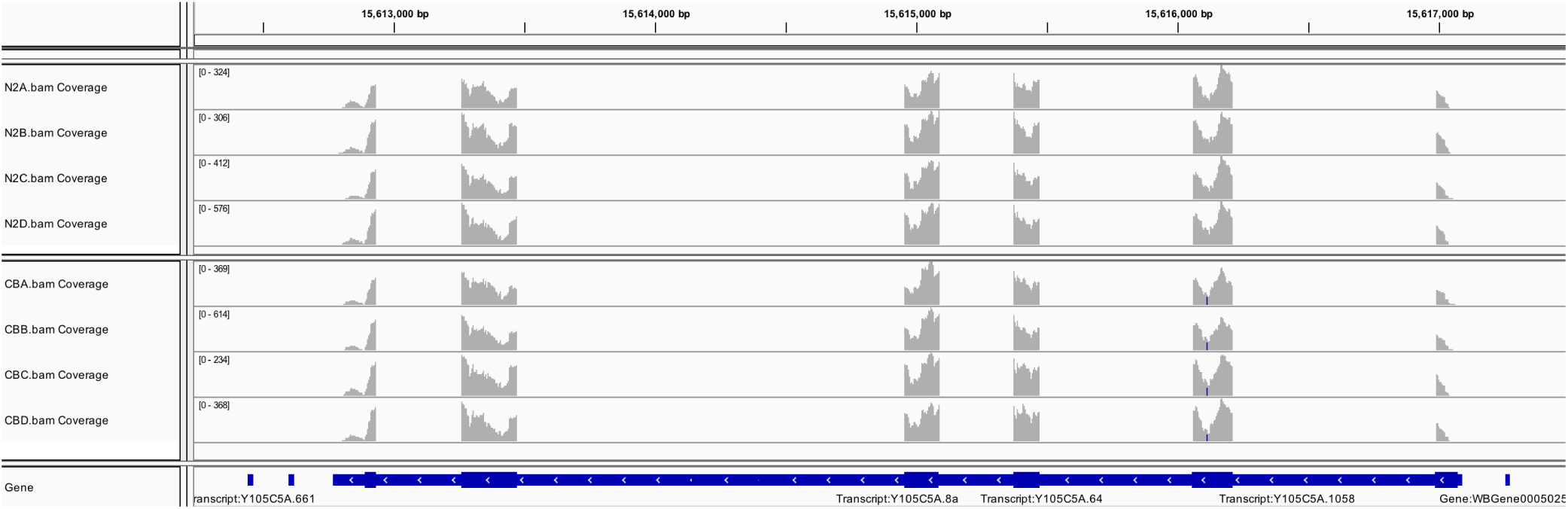
*C. elegans Y105C5A.8* RNA-seq data. RNA-seq alignment coverage shown for four biological replicates of N2 and CB4856.

**S6 Fig.**
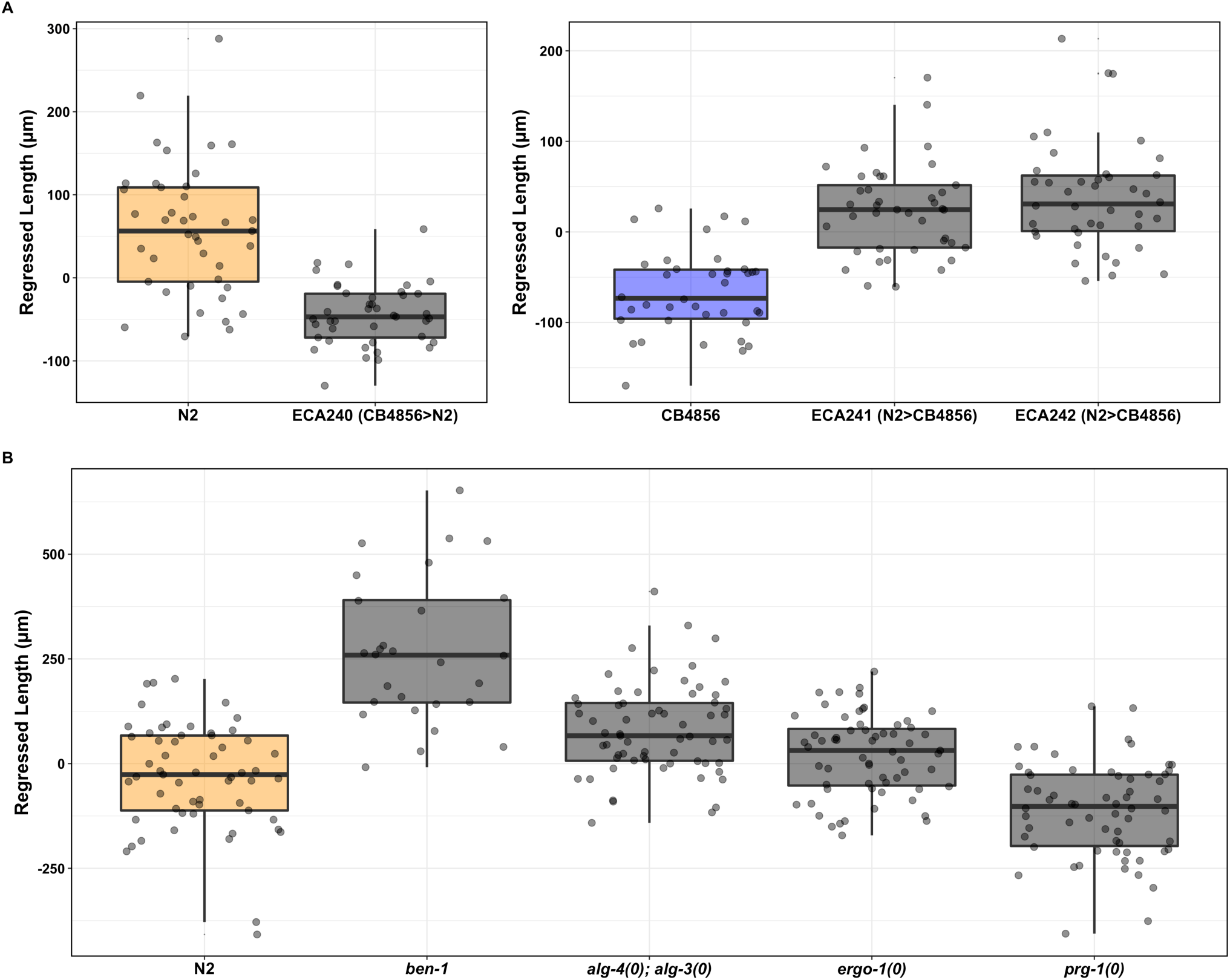
(A) piRNA interval NILs recapitulate QTL direction of effect. Reciprocal NILs covering the primary *C. elegans* piRNA cluster (chromosome IV: 13.5 - 17.2 Mb) were phenotyped in the presence of 12.5 *μ*M albendazole. Tukey box plots show drug effects on animal length (75th quantile shown). The introgression of the piRNA cluster from CB4856 into N2 (ECA240) leads to greater albendazole sensitivity compared to N2 (P < 0.001). Introgression of the piRNA cluster from N2 into CB4856 (ECA241 and ECA242) confers greater albendazole resistance compared to CB4856 (P < 0.001). **(B) Argonaute mutants that converge on the WAGO 22G RNA pathway were phenotyped in 12.5 *μ*M albendazole.** *ergo-1* and *alg-3; alg-4* mutants, which interact with distinct classes of 26G RNAs, do not confer increased albendazole sensitivity phenotype to the N2 genetic background. Loss of *prg-1,* the primary Argonaute associated with 21U-RNA/piRNA activity, confers albendazole sensitivity in the N2 background *(prg-1(0)* vs N2: p < 0.001). However, despite back crossing, the *prg-1(0) (prg-1(n4357))* strain exhibited slower growth rate and reduced brood size throughout assay propagation. A known benzimdazole-resistance allele, *ben-1,* is included as a positive control and basis for comparison of relative effect sizes *(prg-1(0)* vs *ben-1:* p < 0.001). t-tests were used for statistical comparisons (p-values reported in S9 Table).

**S7 Fig.**
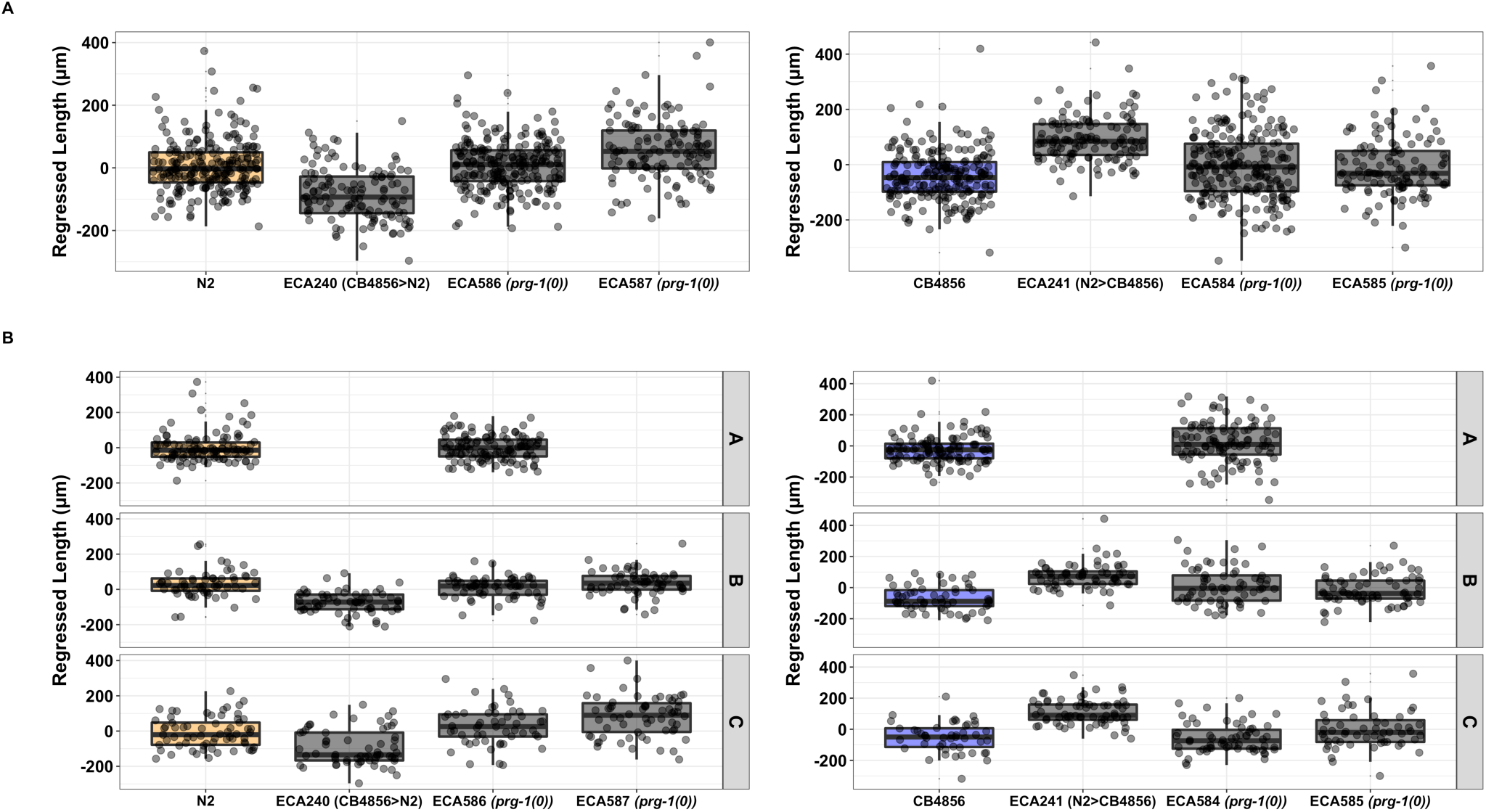
Independent *prg-1* loss-of-function alleles in the N2 and CB4856 backgrounds do not alter albendazole responses. **(A)** Combined data from three independent replicate assays recapitulate the established NIL effect but do not support the hypothesis that this effect is *prg-1* dependent. Tukey box plots show drug effects on animal length (75th quantile shown). Individual assays shown in **(B)** and one-sided t-tests (alpha = 0.05) were used to test whether *prg-1(0)* in the N2 background (ECA586 and ECA587) would lead to greater sensitivity compared to N2 (not significant: p = 0.778 for ECA586; p = 0.9997 for ECA587) and that *prg-1(0)* in the CB4856 background (ECA584 and ECA585) would lead to greater resistance compared to CB4856 (not consistently significant across replicate days). One-tailed t-tests were used for all comparisons (p-values reported in S9 Table).

**S8 Fig.**
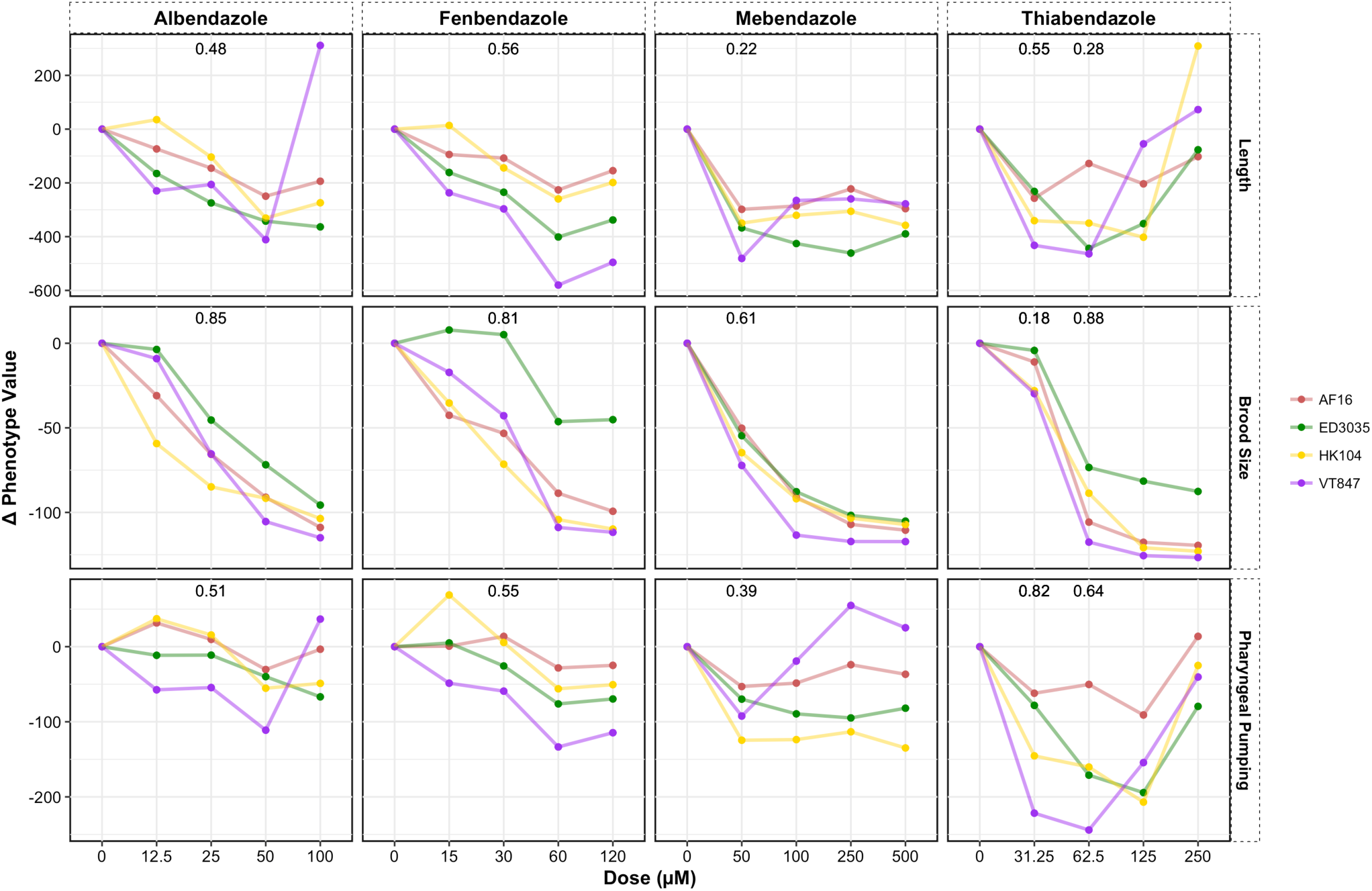
*C. briggsae* benzimidazole dose responses and heritability calculations. Dose responses were carried out with four genetically diverged strains of *C. briggsae.* Phenotypic responses to four drugs are shown with a representative trait for each primary trait group (length, brood size, and pharyngeal pumping). Heritability values are shown for doses used in subsequent linkage mapping experiments. For thiabendazole, we selected a linkage mapping dose (40 *μ*M) that falls between the concentrations where heritability is shown.

**S9 Fig.**
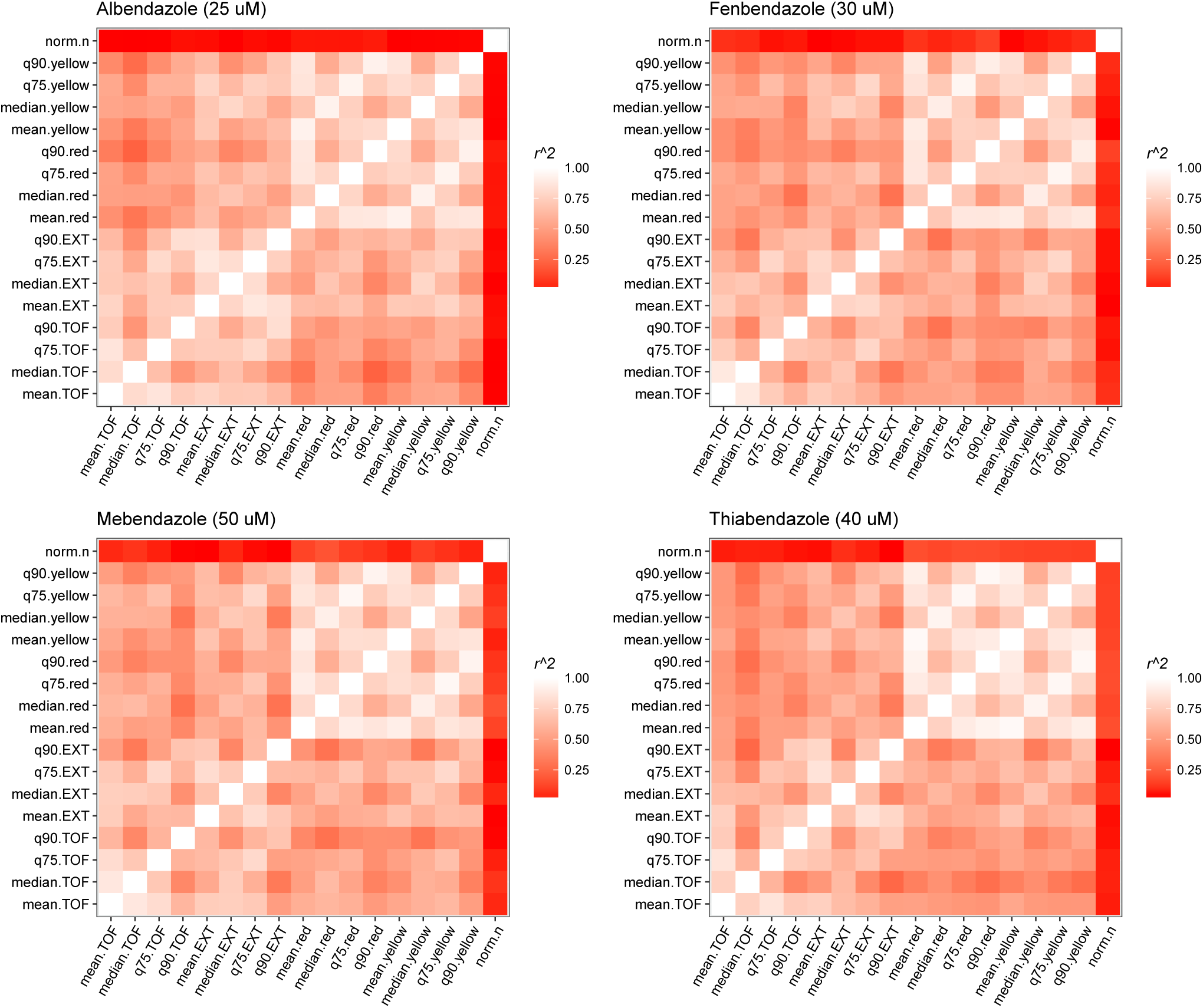
*C. briggsae* summary statistic correlations for primary trait groupings. The correlation structure (Pearson’s correlation coefficient) of summary statistics for measured parameters of animal size (time-of-flight (TOF) and optical density (EXT)), pharyngeal pumping (red and yellow fluorescence), and brood size (norm.n) are shown for each drug tested.

**S10 Fig.**
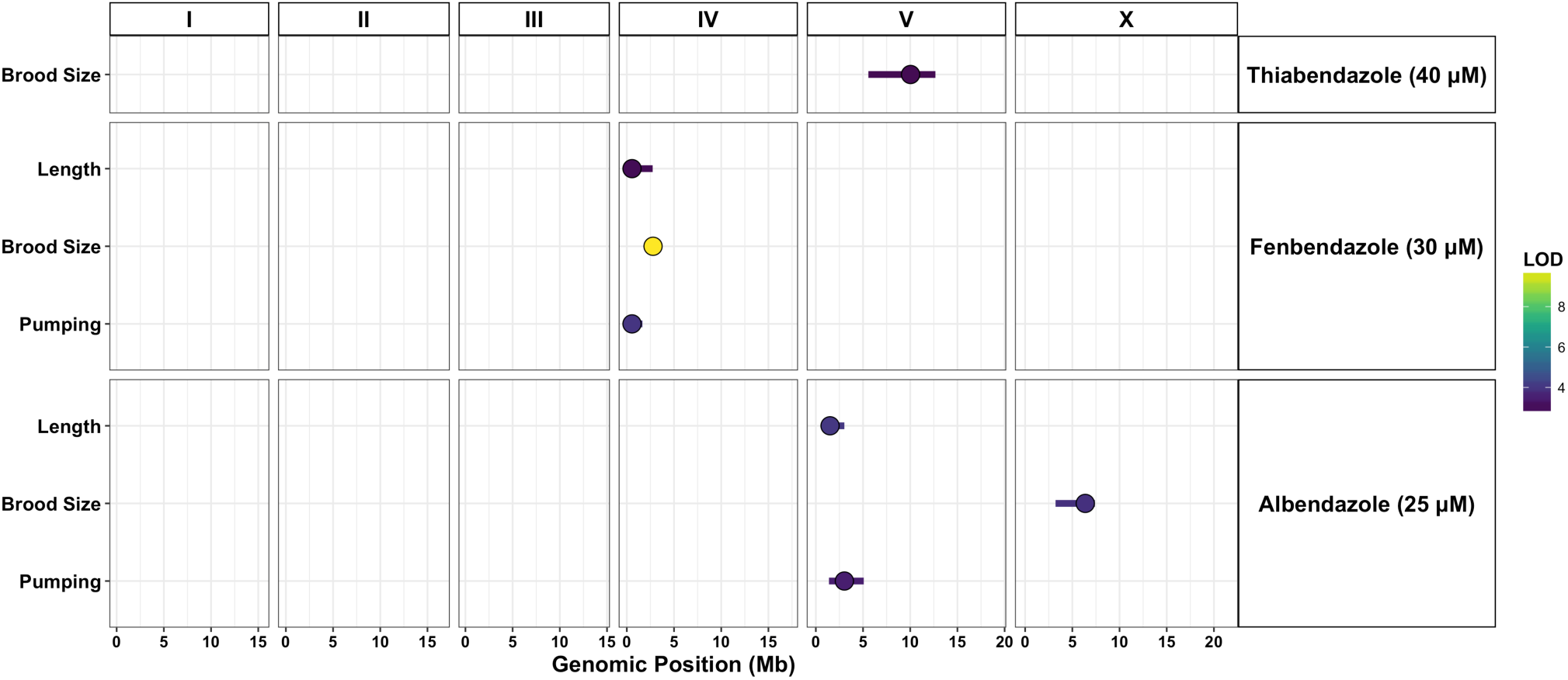
*C. briggsae* benzimidazole QTL grouped by drug and trait. Results of *C. briggsae* linkage mapping experiments are shown for the drug-dose conditions tested and separated by correlated trait group. QTL peak markers (circles) and confidence intervals (lines) are depicted. Fill color corresponds to the QTL LOD score. Overlapping QTL for a given condition-trait group pair are represented by the trait with the highest significance score.

## Supporting Information: Methods

**S1 Methods** Reagents related to NIL and mutant strain assays. Primers and starting RIAILs used in the construction of wholí piRNA-interval NILs. Mutant strains and oligos used to confirm and propagate existing mutant alleles through back-crossing and to construct genome-edited strains.

### *C. elegans* whole piRNA interval NILs

- piRNA interval: Chr IV: 13.5 - 17.2 Mbs / N2 Size: 526, CB4856 Size: 942
- Left Primers (InDel Left: 13207120)
  – oECA904: aacagatactcgccgttgct
  – oECA905: atttgtaccacgcgtgacct
- Right Primers (InDel Right: 17356993) / N2 Size: 443, CB4856 Size: 596
  – oECA910: gacaacgcccactacgacaa
  – oECA911: acccaaccagttgagcacat
- CB4856>N2 NIL (ECA240) / eanIR160 (IV, CB4856>N2) / Starting RIAIL: QX349
- N2>CB4856 NIL (ECA241) / eanIR161 (IV, N2>CB4856) / Starting RIAIL: QX375

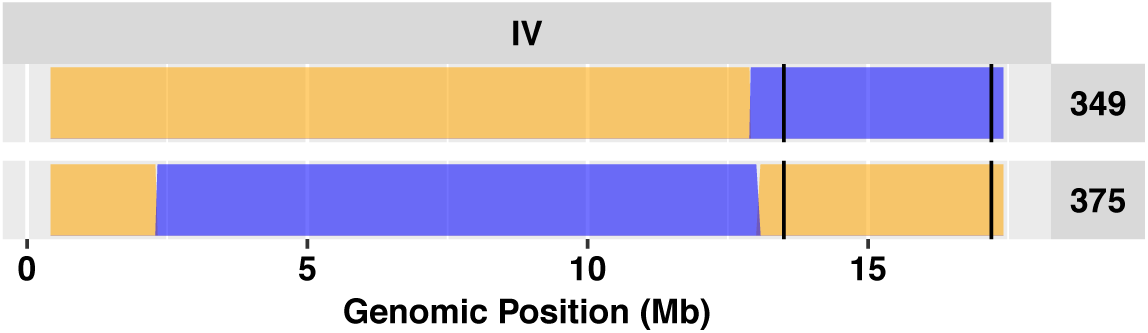

### *C. elegans* mutants

- piRNA mutants confirmed and/or backcrossed in N2 background
  – ECA270: *alg-4(tm1184);alg-3(tm1155)* [ALG-3/4 class 26G RNA]
  – ECA271: WM158 / *ergo-1(tm1860)* [ERGO-1 class 26G RNA]
  – ECA286: *prg-1(n4357)* [21-U RNA]: SX922/ECA269 backcrossed to N2 (9x) and selfed (6x)
  – FX234: *ben-1*
- Mutant confirmation PCR primers
  – *ergo-1(tm1860):* oECA1033 AAGCGTACGAACCCGAGCTT / oECA1034 GAGCGGCTGCTCAGAAGACT
- *prg-1(n4357)* PCR primers for backcrossing
  – oECA1019: AGTCGTGGTACAGATCGTAG
  – oECA1020: GAGAGGCCGTGGTTCAGGAT
  – Wild-type size = 1972, *prg-1(n4-357)* size = 1248 (724 bp deletion)

### *C. elegans* CRISPR *prg-1* loss-of-function strains

- Primers and oligos
  – oECA2002: GUUAGCCUUCGAAUCAACGG
  – oECA2003: CGCUGUGACCGACAAAGCUG
  – oECA2004: GGGTACTATCCAACCCGATCTTTTCATTCG
  – oECA2042: CGCGTTTCGTGACAATGATAAATGCS (3’ wobble to account for variant between N2 and CB4856)
  – *dpy-10* crRNA: GCUACCAUAGGCACCACGAG
  – *dpy-10* repair oligo: CACTTGAACTTCAATACGGCAAGATGAGAATGACTGGAAACCGTACCGCATGCGGT-GCCTA GGTAGCGGAGCTTCACATGGCTTCAGACCAACAGCCTAT
- Strains
  – ECA584: Exon 1-7 deletion (CB4856) and ECA585: Exon 2-7 deletion (CB4856)
  – ECA586: Exon 1-7 deletion (N2) and ECA587: Exon 2-7 deletion (N2)

## Supporting Information: Tables

**S1 Table.** *C. elegans* QTL that explain > 5% of trait variation for all drug-dose combinations tested. Each QTL that was discovered for defined trait groups is annotated with its peak position, confidence interval, estimated variance explained, and effect size.

**S2 Table.** *C. elegans* protein-coding gene variants within the albendazole QTL interval. Previously defined variants distinguishing N2 and CB4856 were annotated based on predicted effects on protein function or expression. Variants of “moderate” or “high” effect are shown for the QTL and the NIL-narrowed interval.

**S3 Table.** *C. elegans* piRNA genes within the albendazole QTL interval. The genomic coordinates, gene IDs, and feature IDs of annotated piRNA genes within the QTL and NIL-narrowed intervals are provided.

**S4 Table.** *C. elegans* N2 and CB4856 young adult RNA-Seq data. Read counts (TPM) and p-values for differentially expressed genes between strains.

**S5 Table.** *C. elegans* expression QTL (eQTL) associated with the NIL-narrowed albendazole QTL interval.

**S6 Table.** *C. briggsae* QTL that explain > 5% of trait variation for all drugs tested. Each QTL that was discovered for defined trait groups is annotated with its peak position, confidence interval, estimated variance explained, and effect size.

**S7 Table.** *C. briggsae* protein-coding gene variants within the fenbendazole QTL interval. Variants distinguishing AF16 and HK104 were annotated based on predicted effects on protein function or expression. Variants of “moderate” or “high” effect are shown for the QTL interval.

**S8 Table.** Parasite *(B. malayi* and *S. ratti)* orthologs of all proteins with variants of “moderate” or “high” impact within *C. elegans* QTL identified.

**S9 Table.** P-values for all statistical tests comparing individual strains (NILs and mutants) with their corresponding parental strains.

